# Cross-Serogroup Analysis of Representative Top-Seven Shiga toxin-producing *Escherichia coli* Plasmids Reveals Lineage-Specific Patterns

**DOI:** 10.64898/2026.09.16.752139

**Authors:** Ali Nemati, Irvin Rivera, Soren Mohammadi, Ali Dadvar, Nima Dehdilani, Sara S. K. Koenig, Adeleh Arzhangi, Anwar Kalalah, Ali Sardarzadeh Majd, Amiryazdan Taghizadeh Shool, Ute Römling, Federica Gigliucci, Mark Eppinger

## Abstract

Plasmids play a significant role in shaping the pathogenic potential of Shiga toxin-producing *Escherichia coli* (STEC) by encoding virulence and adaptive genes that complement chromosomal determinants. While plasmid-encoded factors such as *ehxA*, *espP*, *katP*, and *toxB* are known to enhance intestinal colonization and host interaction, the diversity and evolutionary patterns of STEC plasmids remain insufficiently characterized. Most previous studies have focused on single serogroups, particularly O157, leaving cross-serogroup comparisons largely unexplored. To address this gap, we conducted an in-depth investigation of 109 complete plasmid sequences (1.5–187 kb) from the “Top Seven” STEC serogroups (O26, O45, O103, O111, O121, O145, and O157), retrieved from NCBI, to examine their structural organization, virulence composition, resistance patterns, mobility potential, and evolutionary dynamics. Our analysis revealed the dominance of F-type plasmids carrying IncFIB and IncFII replicons across serogroups, along with lineage-specific associations with major virulence genes. By examining the enterohemolysin operon across *ehxA*-positive plasmids, we identified a highly conserved structural framework maintained across 92.5% of analyzed sequences, with minimal variation. We further characterized antimicrobial resistance gene distribution, finding that only 11% of plasmids (12/109) carried such genes, of which 92% were multidrug-resistant. Analysis of mobility features revealed that predicted conjugative potential and MOBF-type relaxases varied considerably among serogroups. The 32 plasmids without virulence or resistance cargo carried at most colicin determinants, yet all received a predicted mobility class, with conjugative machinery confined to plasmids above 37 kb. At the plasmid level, serogroups O26/O103 are closely related and carry relatively high-risk virulence profiles, while the O121/O145 group exhibits moderate virulence gene inventories. The O45/O111 group is further distinguished by its transfer features, while O157 strains form a distinct clade with the broadest array of plasmid-encoded high-risk virulence genes.

**Importance:** Shiga toxin-producing *E. coli* (STEC) is a globally significant foodborne pathogen responsible for severe gastrointestinal disease and life-threatening complications, particularly in children and vulnerable populations. As our understanding of STEC pathogenicity continues to expand, the question of what constitutes the core plasmid-encoded genetic repertoire across major serogroups is becoming increasingly important. In this study, we revealed that different STEC lineages carry distinct plasmid-borne combinations of disease-promoting, resistance, and transfer-associated genes. Together, the presented findings move beyond single-serogroup perspectives and establish a detailed cross-serogroup framework that can inform future risk assessment, outbreak surveillance, and public health intervention strategies.

**Graphical:** 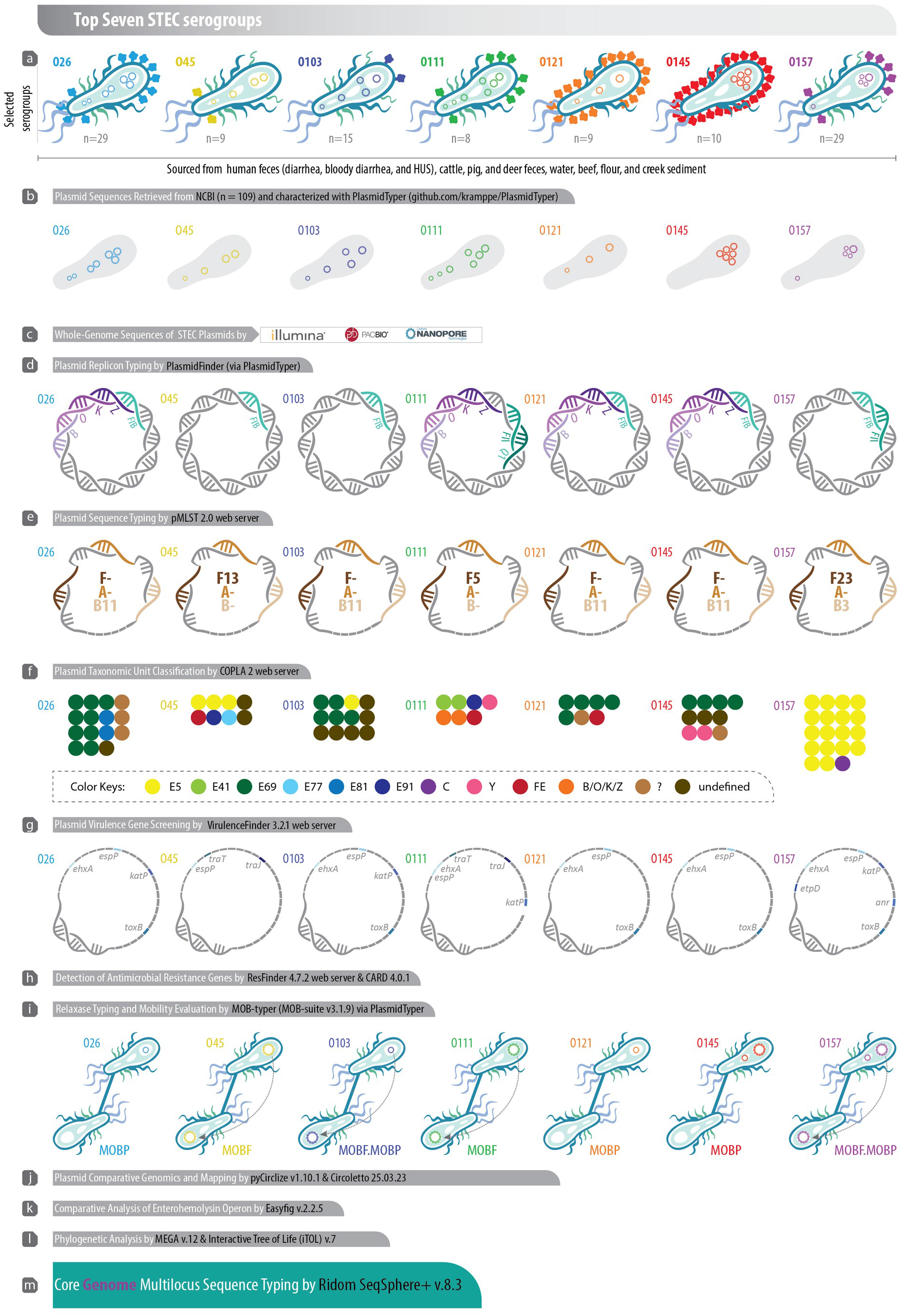

## Background

Plasmids play a vital role in bacterial pathogenesis by functioning as mobile genetic elements that disseminate important traits, including virulence factors, antimicrobial resistance (AMR), and metabolic functions (1). These traits enable bacteria to adapt rapidly to environmental challenges and enhance their survival (2). In Shiga toxin-producing *Escherichia coli* (STEC), plasmids significantly contribute to virulence and disease severity (3–6). While the *stx* genes encoding Shiga toxins are typically phage-borne, and the *eae* gene encoding the adhesion factor intimin is chromosomally encoded, many additional virulence-associated genes are located on large (>90 kb) plasmids (7, 8). These plasmids, especially pO157-like plasmids, are prevalent among major STEC serogroups such as O157, O26, and O111 (9).

Such plasmids often encode factors including *ehxA* (enterohemolysin), *espP* (serine protease), *katP* (catalase-peroxidase), *toxB* (adherence factor), as well as *etpD* and *traT*, which are involved in secretion systems and serum resistance (2, 10). Collectively, these genes promote intestinal colonization, facilitate immune evasion, and enhance bacterial survival under host conditions, ultimately increasing the pathogenic potential of STEC (11). Additionally, a subset of STEC plasmids harbor AMR genes or mobilization elements that enable the horizontal transfer of resistance and other pathogenicity traits between bacteria, contributing to the emergence of more resilient and virulent strains (12).

STEC plasmids vary widely in size, typically ranging from several kilobases (kb) to over 200 kb (2, 13, 14). Large plasmids, often exceeding 90 kb, are of particular interest because they harbor multiple virulence genes, AMR determinants, and conjugation machinery (13). These large plasmids, such as the well-characterized pO157, play a central role in STEC pathogenicity and adaptability (15). Historically, the extraction and sequencing of large plasmids has been challenging; however, dedicated plasmid extraction protocols have substantially improved this process. Short-read sequencing technologies often lack the resolution needed to accurately reconstruct repetitive regions and structural variants, leading to fragmented or incomplete assemblies (16, 17). In contrast, long-read sequencing platforms such as Oxford Nanopore and PacBio have significantly improved the resolution of complex plasmid structures (18, 19). Nevertheless, large plasmids may still be lost during laboratory cultivation under non-selective conditions (20), or integrate into the chromosome. Together, these factors have contributed to the limited availability of complete, well-annotated STEC plasmid sequences, hindering detailed insights into their structure and function.

Despite the recognized importance of plasmids in STEC pathogenicity, comparative analyses of plasmid content across the major STEC serogroups remain limited. Most previous studies of plasmid inventories have focused on individual serogroups, particularly O157 (21, 22), leaving gaps in our understanding of both the shared and unique plasmid features among the broader “Top Seven” STEC. Comparable genome-scale frameworks have recently been established for emerging non-Top Seven serogroups such as O118 (23), underscoring that plasmid-borne virulence gene boundaries differ between STEC lineages. A comprehensive cross-serogroup examination of plasmid content, mobility, and genetic structure is essential to uncover evolutionary patterns, track the dissemination of virulence and resistance determinants, and better understand how these mobile elements shape STEC diversity and adaptability. In this study, we investigated a unique collection of 109 plasmids (1.5–187 kb) associated with the “Top Seven” STEC serogroups (O26, O45, O103, O111, O121, O145, and O157), retrieved from publicly available genomic repositories. Through comprehensive genomic analysis, we aimed to characterize their structural and functional features, assessing replicon types, virulence and AMR genes, plasmid mobility, and inferred phylogenetic relationships. Our findings provide novel insights into the genetic makeup of these plasmids, moving beyond single-serogroup perspectives toward a comprehensive, cross-serogroup framework that contributes valuable information to the growing body of knowledge on STEC plasmid biology and lineage-specific plasmid patterns.

## Methods

### Whole-Genome Sequences of STEC Plasmids

Complete STEC plasmids were retrieved from NCBI in July 2025 by querying the NCBI Assembly and Nucleotide databases using two complementary approaches. First, publications reporting genome assemblies of the “Top Seven” STEC serogroups were identified, and the accession numbers provided by the authors were used to retrieve the corresponding BioProjects, from which associated plasmid sequences were extracted. Second, plasmids were identified directly via a targeted search of the NCBI Nucleotide database, using “Escherichia coli” combined with the O:H serotypes corresponding to the “Top Seven” serogroups; each candidate BioProject was then checked manually to confirm that it included at least one plasmid sequence. Records were retained if they met all of the following criteria: (i) the organism was identified as *E. coli*; (ii) the isolate was annotated as Shiga toxin-producing *E. coli* or carried the relevant *stx* determinant(s); (iii) the genome assembly was complete; (iv) the serogroup was assigned to one of the “Top Seven” serogroups; and (v) the record contained the required metadata, including source host or matrix, geographic location of origin, collection date, sequencing platform, sequence length, and GC content. Duplicate records, incomplete assemblies, draft assemblies, and records lacking a reliable serogroup assignment were excluded. In total, 109 complete plasmid sequences from 61 “Top Seven” STEC strains were retrieved from NCBI. These plasmids were obtained from strains belonging to the serogroups: O26 (n = 29), O45 (n = 9), O103 (n = 15), O111 (n = 8), O121 (n = 9), O145 (n = 10), and O157 (n = 29). The plasmid hosts originated from a wide range of countries globally, including Japan, Belgium, the United Kingdom, the United States, Canada, New Zealand, and Germany, from July 2016 to January 2025 (Tables S2 and S3 in Supplementary File). The strains had been recovered from human feces (diarrhea, bloody diarrhea, and hemolytic uremic syndrome or HUS), cattle, pig, and deer feces, water, beef, flour, and creek sediment (Table S1). Sequences were produced using the PacBio, Illumina MiSeq, and Oxford Nanopore MinION platforms, with sizes ranging from 1,546 to 187,274 base pairs and GC contents between 40% and 54%. For analysis, the 109 plasmids were grouped by cargo. No redundancy filtering was applied; near-identical plasmids carried by different strains are reported as such. The dataset was assembled by querying the NCBI Assembly and Nucleotide databases in July 2025, and retaining all *Escherichia coli* genome assemblies that satisfied the criteria above. Only closed assemblies with a fully resolved plasmid complement were used, so that the entire plasmidome of each strain could be characterized, and no plasmid was excluded on the basis of size, replicon type, or sequence similarity. All plasmids were included in the analyses. The 109 plasmids therefore comprise all qualifying records identified by this search, as a snapshot of the publicly available STEC plasmidome at the time of retrieval, rather than a curated selection. Table S2 lists the 77 plasmids carrying at least one non-bacteriocin virulence gene or at least one acquired resistance gene, and Table S3 lists the 32 plasmids without such cargo (bacteriocin genes only, or no cargo genes detected). Bacteriocin determinants (*cba*, *cea*, *cia*, *cib*, *cma*, *colE2*, *colE8*, *ccI*, *cvaC*, *mchF*) were not regarded as cargo on their own. Prevalence values are given for the whole collection where indicated; serogroup-level profiles (Table 1) refer to the cargo-carrying plasmids of Table S2.

**Table 1.**
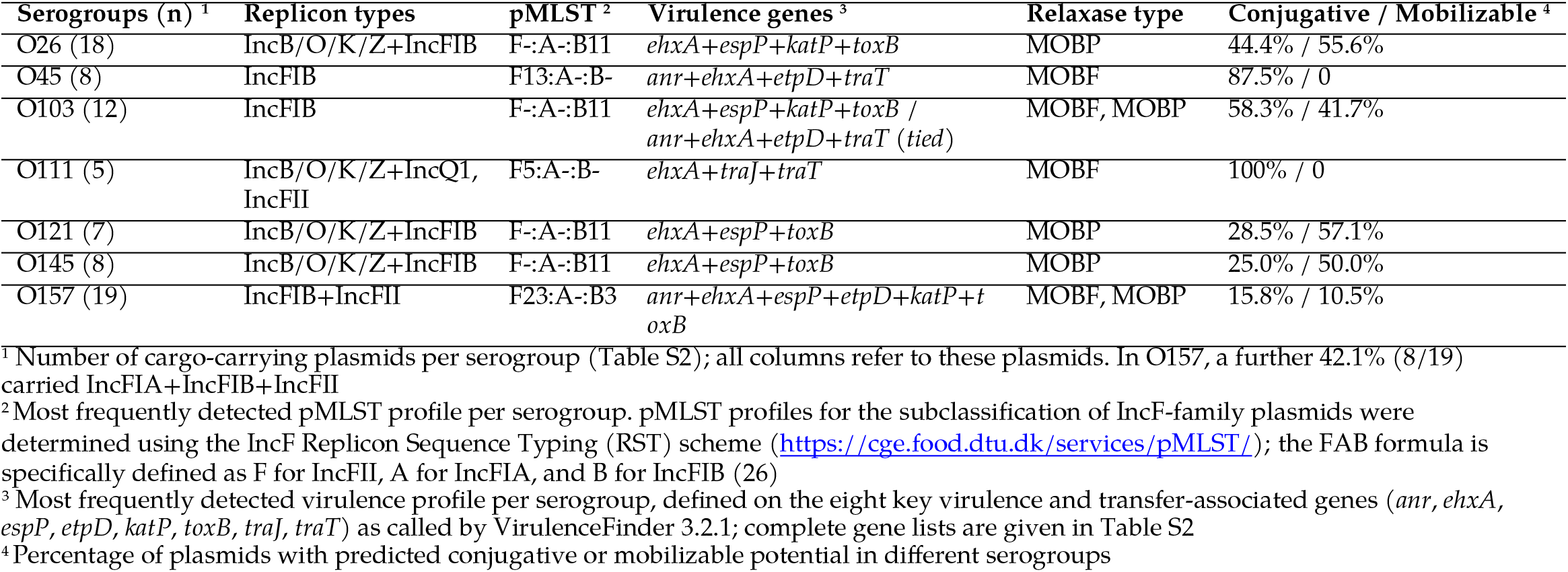
Most frequently detected plasmid profiles and associated features in the “Top Seven” STEC serogroups.

Plasmid sequences were processed and quality-checked using RStudio v2025.05.0-496 running R v4.4.1 and the Biostrings package v2.72.0 (https://bioconductor.org/packages/release/bioc/html/Biostrings.html). Curation involved a multi-step screening pipeline wherein sequence metadata and FASTA headers were first parsed to verify circularized, closed DNA molecules represented as a single contiguous sequence, excluding any records flagged with draft status, incomplete headers, or linear topology without circularity validation. Next, sequence integrity was assessed in Biostrings by analyzing base composition to quantify ambiguous bases (N characters) and internal assembly gaps, removing any records containing >0% ambiguous bases. Finally, cleaned sequences were cross-referenced against the PLSDB web server (https://ccb-microbe.cs.uni-saarland.de/plsdb2025/, accessed 5 July 2025) to confirm plasmid identity, topological status, and alignment with verified STEC plasmid records, ensuring that only sequences meeting all quality thresholds were retained for downstream analysis.

### Plasmid Characterization Workflow

All 109 plasmids were characterized with PlasmidTyper v1.0.1 (https://github.com/kramppe/PlasmidTyper), an open-source workflow developed for this study that submits sequences to, retrieves results from, and collates the output of the typing tools described below (replicon, pMLST, PTU, virulence and resistance genes, relaxase, and mobility). Each tool can be run against its web service or a local installation, so that the analyses can be reproduced offline; discrepancies between the two routes are reported rather than reconciled. Tool versions, access dates, and thresholds are given in the respective sections. The compiled per-plasmid profiles are provided in Tables S2 and S3, and the accession list and compiled profiles are distributed with the repository as an example panel.

### Plasmid Replicon Typing

Plasmid replicon types were identified by analyzing plasmid nucleotide sequences for their replication-associated genes. This was performed using the PlasmidFinder 2.1 web server (https://cge.food.dtu.dk/services/PlasmidFinder/, accessed 7 July 2025). The analyses were performed with default settings, including a minimum sequence identity of 95% and a minimum coverage of 60%. The Enterobacterales database was used for replicon classification (24). Replicon assignments were additionally validated by cross-checking the 29 plasmids present in PLSDB against the PLSDB web server (https://ccb-microbe.cs.uni-saarland.de/plsdb2025/browse, accessed 7 July 2025), yielding consistent results (25). For the final tables, all 109 plasmids were re-typed with a local PlasmidFinder installation (plasmidfinder_db, accessed 2 September 2026) run through PlasmidTyper at the same thresholds. The local run reproduced the web-server calls and additionally resolved IncFIA replicons in eight O157 plasmids and a replicon in five plasmids left untyped by the web server; the local calls are reported in Tables S2 and S3.

### Plasmid Sequence Typing Using pMLST

To achieve higher resolution beyond replicon typing, plasmid multilocus sequence typing (pMLST) was used to characterize the specific sequence types (STs) of the identified Inc-group plasmids (26). This approach identifies STs by analyzing internal fragments of multiple housekeeping genes. The analyses were carried out using the pMLST 2.0 web server (https://cge.food.dtu.dk/services/pMLST/, accessed 26 July 2025). For each plasmid, the corresponding replicon type (Inc group) was specified in the pMLST configuration, and the input sequences were provided in FASTA format (27). For multireplicon plasmids containing both IncF and IncB/O/K/Z elements, subtyping was restricted to the IncF configuration, as no formal pMLST scheme is currently available for the IncB/O/K/Z family on the web server. The broader genetic backgrounds of these hybrid structures were instead evaluated using core backbone gene phylogenetics.

### Computational Plasmid Taxonomic Assignment

Plasmid taxonomic unit (PTU) classification was performed using the COPLA pipeline (Classification of Plasmids) to assign plasmids into phylogenetically defined PTUs based on plasmid sequence similarity and network-based clustering (28). The analyses were conducted using the COPLA 2 web server (https://castillo.dicom.unican.es/copla/, accessed 7 March 2026). Plasmid sequences were uploaded in FASTA format, and the tool classified them by comparing their genomic features, including average nucleotide identity (ANI), gene content similarity, and relaxase type, against a curated reference plasmid database to determine the most probable PTU assignment. The COPLA algorithm integrates hierarchical clustering within plasmid similarity networks to infer PTU membership, enabling standardized classification of plasmids across different incompatibility groups and bacterial hosts (28). If the query plasmid clusters with a previously defined PTU, it is assigned to that PTU; otherwise, it may remain undefined or represent a potential new PTU (PTU-?).

### Plasmid Virulence Gene Screening

Virulence genes present on the plasmids were identified using the VirulenceFinder 3.2.1 genomic tool (https://cge.food.dtu.dk/services/VirulenceFinder/, accessed 3 September 2026). The analyses were conducted by selecting *E. coli* as the target species, with a minimum sequence identity of 90% and a minimum gene coverage of 60%. All remaining parameters were retained at default values (29). Detected plasmid-encoded virulence genes were cross-verified using PLSDB (https://ccb-microbe.cs.uni-saarland.de/plsdb2025/browse, accessed 7 July 2025) (25). All 109 plasmids were screened with VirulenceFinder 3.2.1, and Tables S2, S3 and S5 report these calls; only hits meeting the stated 90% identity and 60% coverage thresholds were counted; the web results page also lists shorter partial alignments below these thresholds, which were excluded. For the summary of virulence profiles per serogroup (Table 1), profiles were defined on the eight plasmid-encoded virulence and transfer-associated genes most frequently detected across the collection (*anr*, *ehxA*, *espP*, *etpD*, *katP*, *toxB*, *traJ* and *traT*); complete gene lists are given in Table S2.

### Detection of Antimicrobial Resistance Genes

Antimicrobial resistance genes encoded on the surveyed STEC plasmids were detected using the ResFinder 4.7.2 web server (http://genepi.food.dtu.dk/resfinder, accessed 7 July 2025). The analyses were performed with a minimum sequence identity threshold of 90% and a minimum length coverage of 60%. The species was selected to *E. coli*, and the input data type was specified as FASTA. All other parameters were kept at their default settings (30). To further validate AMR gene detection, all plasmid sequences were additionally run using the Resistance Gene Identifier (RGI 6.0.5) tool from the Comprehensive Antibiotic Resistance Database (CARD 4.0.1, https://card.mcmaster.ca/home, accessed 19 December 2025), with all parameters set to default (31). The results obtained with CARD were highly consistent with those identified by ResFinder. All AMR gene hits were manually curated to confirm gene identity and annotation accuracy. Plasmids carrying acquired resistance determinants from three or more antimicrobial categories were classified as multidrug-resistant (MDR), following the standardized definitions of Magiorakos et al. (32).

### Relaxase Typing and Mobility Evaluation

The STEC plasmids were analyzed to determine their relaxase type and predicted mobility class. These analyses were carried out for all 109 plasmids with MOB-typer (33), run locally through PlasmidTyper with default parameters, and the calls were cross-checked against the MOB-suite-derived annotations stored in the PLSDB web server (https://ccb-microbe.cs.uni-saarland.de/plsdb2025, accessed 26 July 2025), which agreed for every plasmid for which PLSDB reported a call. Relaxase types were assigned according to the MOB typing scheme based on sequence similarity among relaxase (Mob) proteins, which are responsible for nicking the plasmid DNA at the origin of transfer (oriT). To evaluate plasmid mobility, relaxase detection was complemented by identifying genes encoding the type IV secretion system (T4SS), which is required for the formation of a functional conjugation channel. Plasmids encoding both a MOB region and a T4SS were classified as having predicted conjugative potential, whereas plasmids carrying a MOB region but lacking T4SS components were classified as predicted mobilizable. Plasmids without detectable relaxase genes were classified as predicted non-mobilizable. For the PLSDB cross-check, the “Winner-takes-all strategy” option was enabled, while all other parameters remained at their default values (25, 34).

### Plasmid Comparative Genomics and Mapping

Plasmid comparison maps were generated with pyCirclize v1.10.1 (https://github.com/moshi4/pyCirclize, accessed, 9 September 2026). The plasmid *E. coli* O157:H7 strain Sakai pO157 (Accession No. NC_002128.1) was used as the reference sequence. Each of the 77 cargo-carrying plasmids was aligned against the reference with BLASTn v2.16.0, and the resulting rings were colored by serogroup and shaded by BLASTn nucleotide identity to the reference on a continuous gradient (lighter shading indicates lower identity, spanning 70–100%; regions below 70% identity appear as gaps), with an inner ring displaying the GC skew of the reference sequence. This approach enabled a clear comparison of conserved and variable regions among the plasmids (35).

Comparative plasmid sequence similarity analyses were performed using Circoletto v25.03.23 (https://bat.infspire.org/circoletto/, accessed 1 January 2026), a Circos-based visualization tool that displays pairwise BLASTn alignments as ribbons between circular ideograms (36). Due to the high virulence gene content and epidemiological relevance of *E. coli* O157, plasmids from this serogroup were used as the reference framework. To represent the genetic diversity of the plasmid population, three plasmids from each non-O157 serogroup (O26, O45, O103, O111, O121, and O145) were chosen based on size, including the smallest, median-sized, and largest available sequences. These plasmids were compared pairwise against the *E. coli* O157:H7 str. Sakai pO157 reference plasmid (Accession No. NC_002128.1) and one additional large O157 plasmid from our dataset (Accession No. CP043023.1), which originated from an HUS case. BLASTn was performed with an E-value threshold of 1e−10, and only the best hit per query was retained. Ribbons were colored based on the relative BLAST bit score using a score/max ratio, where blue represents low similarity (≤0.25 of the maximum score), green indicates moderate similarity (>0.25–0.50), orange reflects high similarity (>0.50–0.75), and red denotes very high similarity (>0.75). This relative color scaling emphasizes the strongest homologous regions within each comparison while accounting for differences in plasmid size and gene content.

### Comparative Analysis of Enterohemolysin Operon

Considering the key role of hemolysin in the pathogenicity of STEC and the notable presence of hemolysin genes among the plasmids analyzed, the genetic organization of the enterohemolysin (*ehx*) operon was further investigated. Plasmids harboring the complete *ehxCABD* operon (n = 40) were selected for comparative analysis. The structural arrangement and gene homology within the operon were first examined using Easyfig v2.2.5 (https://mjsull.github.io/Easyfig/, accessed 5 October 2025), which enables visualization of genomic regions and BLAST-based comparisons among multiple plasmids (37). Default parameters were used to generate pairwise comparisons and gene annotations. Subsequently, representative protein sequences (EhxA, EhxB, EhxC, and EhxD) from these operons, including those exhibiting divergent gene contexts or lower sequence similarity, were aligned with reference sequences using ClustalW (https://www.genome.jp/tools-bin/clustalw, accessed 24 February 2026) (38). Specifically, proteins from the pO157 plasmid of *E. coli* O157:H7 str. Sakai (Accession No. NC_002128.1) and the alpha-hemolysin operons of uropathogenic *E. coli* 536 (Accession No. NC_008253.1) were included as anchor points for the analysis. Additionally, the EhxA and EhxB sequences from the *E. coli* plasmid pHly152 (Accession No. M14107.1) were incorporated to provide a comprehensive evolutionary framework covering both enterohemolysin and alpha-hemolysin clades. Phylogenetic reconstruction was performed in MEGA v12 (https://megasoftware.net/, accessed 24 February 2026), using the Neighbor-Joining method with 1,000 bootstrap replicates (39).

### Phylogenetic Analysis

To explore the relatedness of plasmid backbones and their association with mobility-related features (relaxase type) and key virulence genes (*ehxA*, *toxB*, *espP*, and *katP*), a phylogenetic analysis was conducted using three conserved plasmid-associated genes (*aidA*, *repB*, and *finO*). These genes were selected as stable phylogenetic markers due to high prevalence and conserved nature across the analyzed plasmids, as identified via pangenome examination using Roary v3.13.0 (minimum BLASTP identity 90%) from the GFF3 files produced by Prokka v1.14.6 annotation of all 109 plasmids (Table S4 of the Supplementary File; https://sanger-pathogens.github.io/Roary/, accessed 5 October 2025) (40). The three genes were aligned individually with MAFFT v7 and concatenated into a supermatrix (71 plasmids carrying at least one of the three genes; 2,493 aligned positions), which was analyzed in MEGA v12 (https://megasoftware.net/, accessed 5 October 2025). The optimal nucleotide substitution model was selected based on the lowest Bayesian Information Criterion (BIC) score (Kimura 2-parameter with gamma-distributed rates and a proportion of invariant sites; K2+G+I). Phylogenetic reconstruction was performed using the Maximum Likelihood (ML) approach with adaptive bootstrapping to evaluate node robustness (39). The resulting phylogenetic tree was visualized and annotated in the Interactive Tree of Life (iTOL) v7 (https://itol.embl.de/, accessed 5 October 2025), incorporating relevant metadata, including serogroup, relaxase type, mobility, and key virulence genes, to enable a comprehensive interpretation of the plasmid lineages (41).

To place plasmid diversity in a broader genomic context and to evaluate relationships among strains beyond serogroup designation, a core genome multilocus sequence typing (cgMLST) analysis was performed based on chromosomal sequences. The closed chromosomes of the “Top Seven” STEC strains from which the analyzed plasmids were derived (Table S1 in Supplementary File), along with the sequence of K-12 substrain MG1655 (Accession No. U00096.3) (42), were imported into Ridom SeqSphere+ v8.3 (Ridom GmbH, Münster, Germany) for core genome (cg) and targeted Multilocus Sequence Typing (MLST) (43–45). The Sequence Type (ST) was determined according to the EnteroBase schema (45). Allele sequences for the Achtman scheme, targeting seven housekeeping genes (*adk*, *fumC*, *gyrB*, *icd*, *mdh*, *purA*, and *recA*), were accessed on the EnteroBase website (https://enterobase.warwick.ac.uk/species/ecoli/download_7_gene). A cgMLST schema was developed using the closed chromosome of K-12 substrain MG1655 (42) as the seed, as previously described (46). Core and accessory MLST targets were identified according to the inclusion/exclusion criteria of the SeqSphere+ Target Definer. Allele information from the targeted seven-gene schema and the defined core genome genes of the panel strains were used to establish phylogenetic hypotheses using the minimum-spanning method with default settings (47, 48).

### Statistical Analysis

Associations between categorical plasmid features (relaxase family, cargo type, serogroup, and predicted mobility class) were tested with two-sided Fisher’s exact tests across all 109 plasmids using SciPy v1.17 (scipy.stats.fisher_exact). The association between serogroup and plasmid backbone similarity to the pO157 reference among the 77 cargo-carrying plasmids compared in Fig. 2 was additionally assessed with a chi-square test of independence (scipy.stats.chi2_contingency); P < 0.05 was considered significant. Counts underlying each test are given in the Results.

## Results

### Compiled Plasmid Profiles

Processing the 109 plasmids through PlasmidTyper yielded one standardized profile per plasmid, combining taxonomy (replicon type, pMLST, PTU), mobility (relaxase type, MPF type, predicted mobility class) and gene inventory (virulence and resistance genes) with the sequence metadata (Tables S2 and S3 in Supplementary File). Across the collection, a replicon type was assigned to 79 plasmids (72.5%), a defined PTU to 82 (75.2%; a further 5 represent potential new PTUs), a relaxase type to 79 (72.5%) and a predicted mobility class to all 109 (40 conjugative, 42 mobilizable, 27 non-mobilizable). Virulence-associated genes were detected in 84 plasmids (77.1%), 74 of which carried at least one gene other than a bacteriocin determinant, and acquired resistance genes in 12 (11.0%). The 77 plasmids carrying virulence or resistance cargo (Table S2) are the subject of the serogroup-level analyses below; the 32 plasmids without such cargo (Table S3) are described alongside each feature.

### Replicon Types

Plasmid replicon typing showed that IncFIB and IncFII replicons, either alone or in combination with other types, were the most prevalent across the entire plasmid collection, highlighting their widespread distribution among the “Top Seven” STEC serogroups (Table 1 and Fig. 1). At the serogroup level, the dominant replicon profile among O26 isolates was IncB/O/K/Z+IncFIB, detected in 44.4% (8/18) of plasmids. In the O45 serogroup, IncFIB was the most frequent replicon, identified in 37.5% (3/8) of plasmids. Among O103 strains, IncFIB was the most common replicon, present in 50.0% (6/12). In O111, IncB/O/K/Z+IncQ1 and IncFII replicons were each detected in 40.0% (2/5) of plasmids. The dominant replicon profile in both O121 and O145 serogroups was IncB/O/K/Z+IncFIB, found in 71.4% (5/7) and 50.0% (4/8) of plasmids, respectively. In O157 strains, IncF multireplicon plasmids accounted for 94.7% (18/19) of plasmids, as IncFIB+IncFII in 52.6% (10/19) and as IncFIA+IncFIB+IncFII in 42.1% (8/19); no O157 plasmid carried an IncB/O/K/Z replicon. Among the 32 plasmids without cargo (Table S3 in Supplementary File), a recognized replicon was identified in only seven (IncY in two O145 plasmids; Col(MG828), Col156, pEC4115, IncI2 and p0111 in one plasmid each), whereas the remaining 25 carried no replicon detectable by PlasmidFinder.

**Fig 1.**
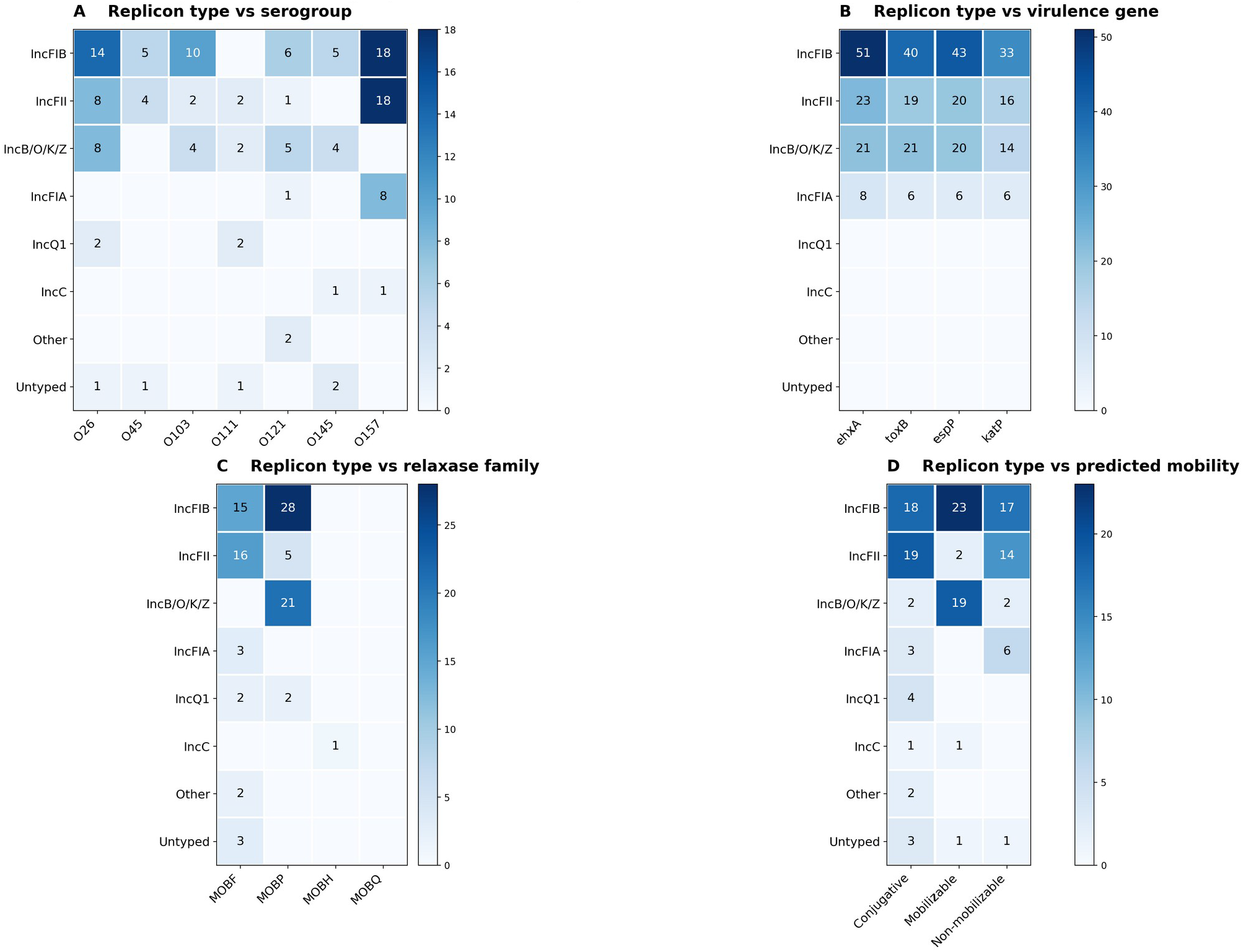
Association of plasmid replicon types with (A) serogroup, (B) the four major virulence genes, (C) relaxase family, and (D) predicted mobility among the 77 cargo-carrying “Top Seven” STEC plasmids (Table S2). Cell values are plasmid counts; color is scaled within each panel. Plasmids with multiple replicons contribute to each replicon row (“Other” = IncFIC, IncX1).

### Subclassification of IncF Replicons via pMLST

pMLST analysis identified the F-:A-:B11 sequence type as the most frequently detected IncF subclassification across multiple serogroups, including O26, O103, O121, and O145, highlighting its widespread distribution and potential importance in the dissemination and maintenance of plasmid-associated traits among “Top Seven” STEC plasmids (Table 1). In O26 strains, F-:A-:B11 was detected in 50.0% (9/18) of plasmids; in 41.7% (5/12) of O103 plasmids, 57.1% (4/7) of O121 plasmids, and 50.0% (4/8) of O145 plasmids. In O45 strains, the dominant sequence type was F13:A-:B-, identified in 37.5% (3/8) of plasmids. For the O111 serogroup, only two plasmids had defined sequence types, both identified as F5:A-:B- (40.0%, 2/5). Finally, plasmids from O157 strains were most commonly assigned to the F23:A-:B3 sequence type, detected in 52.6% (10/19) of cases. Interestingly, the IncF sequence types F13:A-:B-, F5:A-:B-, and F23:A-:B3 were exclusively detected in O45, O111, and O157 serogroups, respectively, suggesting possible serogroup-specific plasmid lineages or evolutionary adaptation (Table 1). Three further plasmids (6.2–49.7 kb) carried an IncF replicon and were typed as F59:A-:B- and F103:A-:B- (two O26 plasmids) and F-:A-:B- (the O45 plasmid CP051654.1, reflecting an imperfect FIB allele match (26)); none of the plasmids without cargo (Table S3 in Supplementary File) carried an IncF replicon.

### PTU Diversity Across STEC Serogroups

COPLA analysis of the 77 cargo-carrying plasmids revealed several PTUs, with PTU-E69 and PTU-E5 representing the most common plasmid lineages identified in this dataset (Table S2 in Supplementary File). PTU-E69 was the most prevalent backbone in plasmids from O26 (50.0%, 9/18), O103 (41.7%, 5/12), O121 (71.4%, 5/7), and O145 (50.0%, 4/8) serogroups. In contrast, PTU-E5 was the primary lineage for O45 (37.5%, 3/8) and O157 (94.7%, 18/19) isolates. The O111 serogroup displayed a more diverse distribution, characterized by PTU-E41 (40.0%, 2/5) and PTU-B/O/K/Z (40.0%, 2/5). Other minor PTUs, including PTU-E81, PTU-FE, PTU-E77, and PTU-C, were sporadically detected across different serogroups (see Graphical Abstract and Table S2 in Supplementary File). Among the 32 plasmids without cargo, several lineages recurred across serogroups: PTU-E14 (n = 6; 6.6–6.7 kb; O26, O103 and O157), PTU-E28 (n = 5; 37.6–38.8 kb; O157, O111 and O121), PTU-E20 (n = 3; 6.8–10.9 kb; O26 and O103) and PTU-E87 (n = 3; 2.7–3.8 kb; O157), while PTU-E91 (O45 and O111) comprised two plasmids and PTU-Y (O145 and O111) comprised three plasmids, both 79–96 kb; two further PTUs (PTU-E241 and PTU-I2) were each represented by a single plasmid, and eight plasmids remained unassigned (Table S3 in Supplementary File). Three O26 plasmids of identical size (6,715 bp; AP027169.1, AP027174.1 and AP027200.1) shared identical relaxase profiles and most likely represent the same plasmid carried by three strains.

### Plasmid-Encoded Virulence Gene Profiles

Screening of plasmid-encoded virulence genes showed that *ehxA* encoding hemolysin was the most consistently detected virulence factor across all serogroups, present in 68.8% (53/77) of cargo-carrying plasmids (48.6%, 53/109, of the whole collection); highlighting its widespread distribution and potential importance in the pathogenicity of “Top Seven” STEC plasmids. This was followed in prevalence by espP (55.8%, 43/77), toxB (51.9%, 40/77), and katP (42.9%, 33/77); anr (42.9%, 33/77) was likewise detected in more than 40% of plasmids (Table 1 and Supplementary Table S5). Across individual serogroups (profiles defined on the eight key genes, see Methods), the most common virulence profile in O26 strains was *ehxA*+*espP*+*katP*+*toxB*, detected in 44.4% (8/18) of plasmids. Among O45 strains, no single profile predominated; anr+ehxA+etpD+traT was the most frequent (2/8), while espP+traJ+traT fell to 1/8 and traT alone was present in 87.5% (7/8) of plasmids. In the O103 serogroup, ehxA+espP+katP+toxB was tied with anr+ehxA+etpD+traT as the most frequent profile, each identified in 33.3% (4/12) of strains. The dominant profile in O111 strains was ehxA+traJ+traT, present in 40.0% (2/5) of plasmids. In O121 strains, *ehxA*+*espP*+*toxB* was the most commonly detected profile (71.4%, 5/7), while in O145 strains the same profile was identified in 25.0% (2/8) of plasmids, whereas a further 25.0% (2/8) of O145 plasmids carried none of the eight key genes. For the O157 serogroup, the most frequent profile was *anr*+*ehxA*+*espP*+*etpD*+*katP*+*toxB*, detected in 84.2% (16/19) of strains. Four plasmids of 6.2–49.7 kb qualified for Table S2 on cargo: CP027332.1 and AP027199.1 (O26; *traT*, and *traT*+*anr*), CP051654.1 (O45, 45,063 bp), which carries a recognizable afaA–D+espP virulence module, and AP027175.1 (O26), which carries resistance genes only. Conversely, five large plasmids (78–96 kb; CP031351.1 and CP027320.1 [O145], CP031917.1 [O45] and AP018804.1 and AP018798.1 [O111]) carried no virulence or resistance genes and are grouped in Table S3, underscoring that classification followed gene content rather than plasmid size. The 32 plasmids without cargo carried at most bacteriocin determinants (cba, n = 7; colE2, n = 3; colE8, n = 1; 10 plasmids), and the *ehxA*/*espP*/*katP*/*toxB* module was entirely absent from them (Table S3 in Supplementary File).

### Plasmid-Encoded Antimicrobial Resistance Gene Profiles

Antimicrobial resistance (AMR) genes were identified in 11.0% (12/109) of all plasmids, corresponding to 15.6% (12/77) of the cargo-carrying plasmids of Table S2, with 91.7% (11/12) of these meeting the criteria for multidrug resistance (MDR). The highest prevalence of AMR-positive plasmids was observed in the O26 (n = 5) and O111 (n = 3) serogroups. Across the collection, resistance to sulfonamides (*sul1*, *sul2*), aminoglycosides (*aph(3’’)-Ib*, *aph*(*6*)*-Id*), and tetracyclines (*tet(A)*, *tet(C)*) represented the most frequent AMR profiles. In contrast, genes associated with resistance to critically important antibiotics, such as colistin (*mcr-5.1*) and macrolides (*mph(A)*), were detected exclusively in the O111 serogroup, representing the least frequent markers. Detailed AMR gene compositions for each serogroup, including sporadically detected genes like *bla*_TEM-1B_ and *dfrA* variants, are provided in Supplementary Table S2; notably, no AMR genes were detected in O45-associated plasmids. Cargo type tracked closely with predicted mobility. All 12 AMR-carrying plasmids were transfer-competent (10 conjugative, 2 mobilizable; 12/12 versus 70/97 of the plasmids without AMR genes, Fisher’s exact test, P = 0.036), and every one of the 30 plasmids carrying the transfer/serum-resistance genes *traT* or *traJ* was predicted conjugative (30/30 versus 10/79, P = 1.4 × 10⁻¹⁸). The pO157-type virulence module (*ehxA*+*espP*+*katP*+*toxB*), by contrast, was found exclusively on mobilizable or non-mobilizable plasmids (15 and 14, respectively) and never on a conjugative one (0/29 versus 40/80, P = 1.5 × 10⁻⁷), indicating that the resident virulence plasmids and the transfer- and resistance-associated plasmids occupy distinct mobility classes.

### Relaxase Types and Conjugation Potential

A notable trend was observed regarding plasmid mobility and relaxase types. Plasmids carrying the MOBF relaxase type were most frequently associated with predicted conjugative potential (hereafter PCP), with all 28 MOBF-typed cargo-carrying plasmids (100%) classified as having PCP, whereas plasmids carrying MOBP alone were predominantly predicted mobilizable (82.1%, 23/28). Across all 109 plasmids, MOBF carriage was strongly associated with predicted conjugative potential (28/31 versus 12/78 for other or undefined relaxases; Fisher’s exact test, P = 1.9 × 10⁻¹³). Notably, the O45 serogroup had the highest proportion of plasmids with PCP, whereas O157 harbored the largest fraction of predicted non-mobilizable plasmids (Table 1). Analysis of relaxase types revealed that MOBP was the most prevalent among O26 plasmids, identified in 66.7% (12/18) of cases. Of the 18 O26-associated plasmids, 8 (44.4%) were classified as having PCP. In O45 strains, MOBF was the dominant relaxase type, present in 87.5% (7/8) of plasmids, and 7 of 8 plasmids (87.5%) showed PCP. Among O103 strains, MOBF and MOBP were each detected in 41.7% (5/12) of plasmids and both in a further two, and seven plasmids (58.3%) were predicted to have conjugative potential. In the O111 serogroup, MOBF was the most common type (60.0%, 3/5), with five plasmids (100%) classified as having PCP. For O121 strains, MOBP was detected in 57.1% (4/7) of plasmids, while two plasmids (28.5%) showed PCP. In O145 strains, MOBP was detected in 37.5% (3/8) and MOBF in 25.0% (2/8) of plasmids, three plasmids (37.5%) lacked a defined relaxase type, and only two plasmids (25.0%) were classified as having PCP. In O157 strains, most plasmids (73.7%, 14/19) did not have a defined relaxase, while MOBF and MOBP were each identified in 10.5% (2/19) of plasmids and MOBH in one. Only three plasmids (15.8%) in this serogroup were classified as having PCP. In contrast to replicon typing, relaxase-based typing resolved the plasmids without cargo: a predicted mobility class was assigned to all 32 (17 mobilizable, 6 conjugative and 9 non-mobilizable), including the 25 without a detectable replicon; MOBP was the most common relaxase family (16/32), followed by MOBF and MOBQ (3/32 each), while 10 lacked a detectable relaxase, one of which (AP038986.1, 1,546 bp) was nevertheless predicted mobilizable on the basis of an origin of transfer (oriT) alone. Across the whole collection, plasmid size and predicted mobility were sharply partitioned: no conjugative plasmid was smaller than 20.8 kb, whereas all plasmids ≤11 kb except one (CP038370.1, 2,731 bp, non-mobilizable) were predicted mobilizable, i.e., they encode a relaxase or oriT but no mating-pair formation system and depend on co-resident conjugative plasmids for transfer (Table S3 in Supplementary File).

### Comparative Genomic Features of STEC Plasmids

Comparative sequence analysis of the plasmids revealed that those associated with O157 strains encoded a higher number of genes and a more diverse functional gene content compared with plasmids from other serogroups (Fig. 2). Notably, the O157 plasmids, which exist in different variants (21, 22) harbored a broader repertoire of genes associated with secretion and transport systems, including components of the type II secretion (*etp*/*gsp*) machinery (49, 50). In contrast, these genes were partially or entirely absent in plasmids from non-O157 serogroups (Fig. 2). Furthermore, hemolysin-associated genes, including *ehxC, ehxA, ehxB, and ehxD*, were detected in most of the “Top Seven” STEC plasmids analyzed, indicating the widespread prevalence of the enterohemolysin operon among major pathogenic STEC lineages (Fig. 2). A notable observation was the consistent presence of the *sopB* gene in the O157 plasmids (Fig. 2). In F-type plasmids such as pO157, *sopB* forms part of the *sopABC* partitioning system, which ensures accurate plasmid segregation during cell division (51). Its frequent occurrence suggests the importance of plasmid stability mechanisms in maintaining and propagating O157 virulence plasmids. Overall, backbone similarity to the pO157 reference was strongly serogroup-dependent: 18 of 19 O157 plasmids shared ≥50% of the reference at ≥70% nucleotide identity, versus only 17 of 58 non-O157 plasmids (χ² = 40.6, df = 6, P = 3.4 × 10⁻⁷ across serogroups; O157 vs non-O157, Fisher’s exact P = 3.8 × 10⁻⁷).

**Fig 2.**
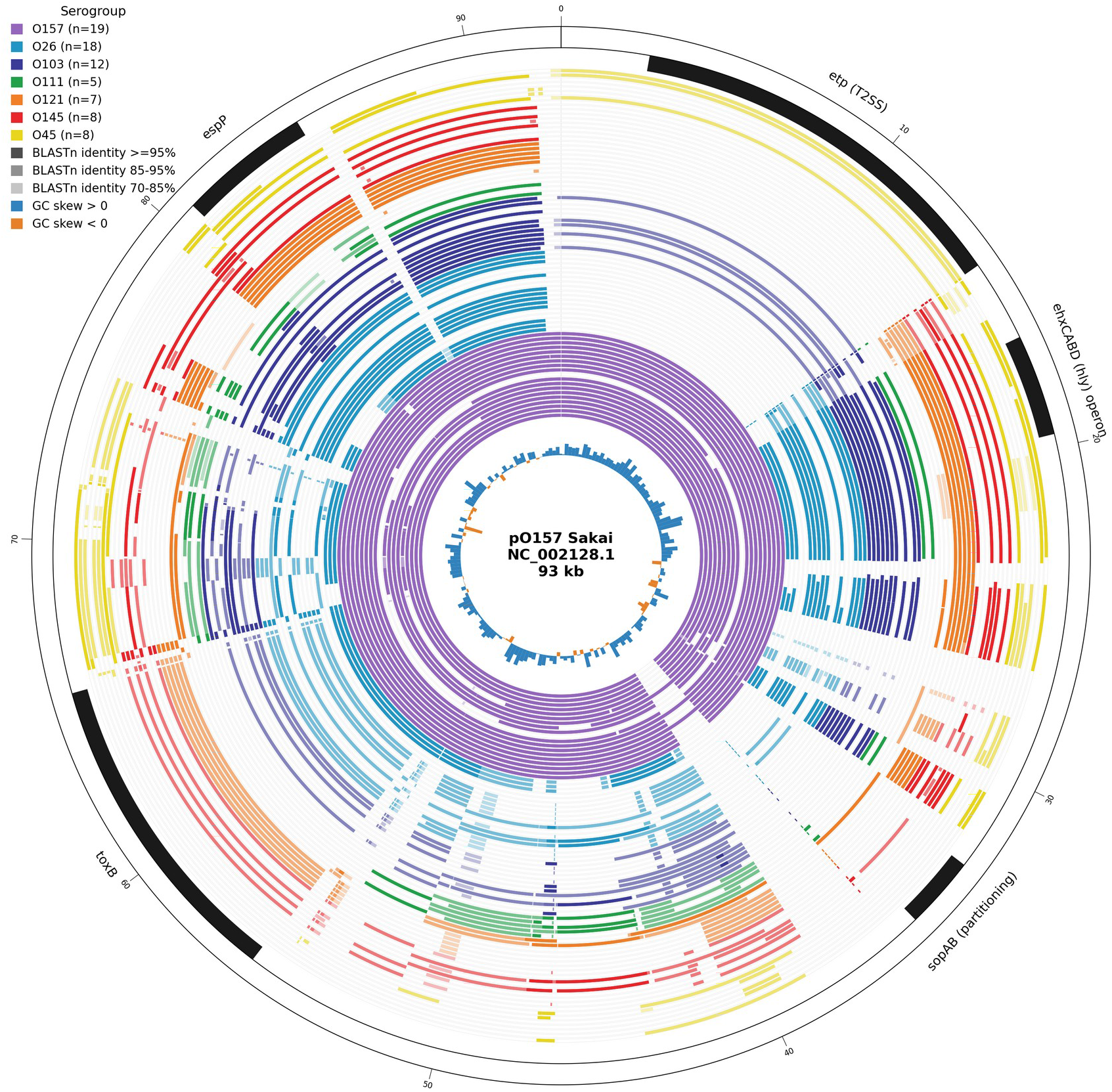
Circular whole-plasmid comparison of the 77 cargo-carrying STEC plasmids (Table S2), colored by serogroup, aligned with BLASTn against *E. coli* O157:H7 str. Sakai pO157 (NC_002128.1) and visualized with pyCirclize; rings are shaded by BLASTn nucleotide identity to the reference on a continuous gradient (lighter shading indicates lower identity, 70–100%; regions absent or below 70% identity appear as gaps), an inner ring shows the reference GC skew (blue, positive strand; orange, negative), and the outer track (black) marks the reference virulence and maintenance loci: the *etp* type II secretion cluster, the *ehxCABD* (*hly*) enterohemolysin operon, the *sopAB* partitioning region, *toxB*, *katP*, and *espP*. Regions absent from a plasmid appear as gaps in its ring.

### Circoletto-based plasmid comparison

Comparative genomic analysis across all serogroups (Fig. 3, A–F) revealed that a notable portion of shared plasmid regions exhibited low-to-moderate sequence similarity (blue ribbons: score ≤0.25; green ribbons: score >0.25–0.50). These conserved segments correspond to essential backbone elements, such as replication and maintenance regions. In contrast, highly conserved regions with alignment scores >0.75 (red ribbons) were more restricted and unevenly distributed, suggesting that vertical inheritance of these specific loci is limited. Notably, plasmids from the O26 and O103 serogroups showed a higher degree of similarity to O157 plasmids compared to other serogroups. Furthermore, the internal comparison of O157 plasmids (Fig. 3, G) demonstrated strong conservation of the core backbone regardless of plasmid size. Overall, these results support a model in which O157 plasmids maintain a highly conserved structural framework, while non-O157 plasmids exhibit variable degrees of relatedness, likely driven by the differential acquisition and loss of accessory regions.

**Fig 3.**
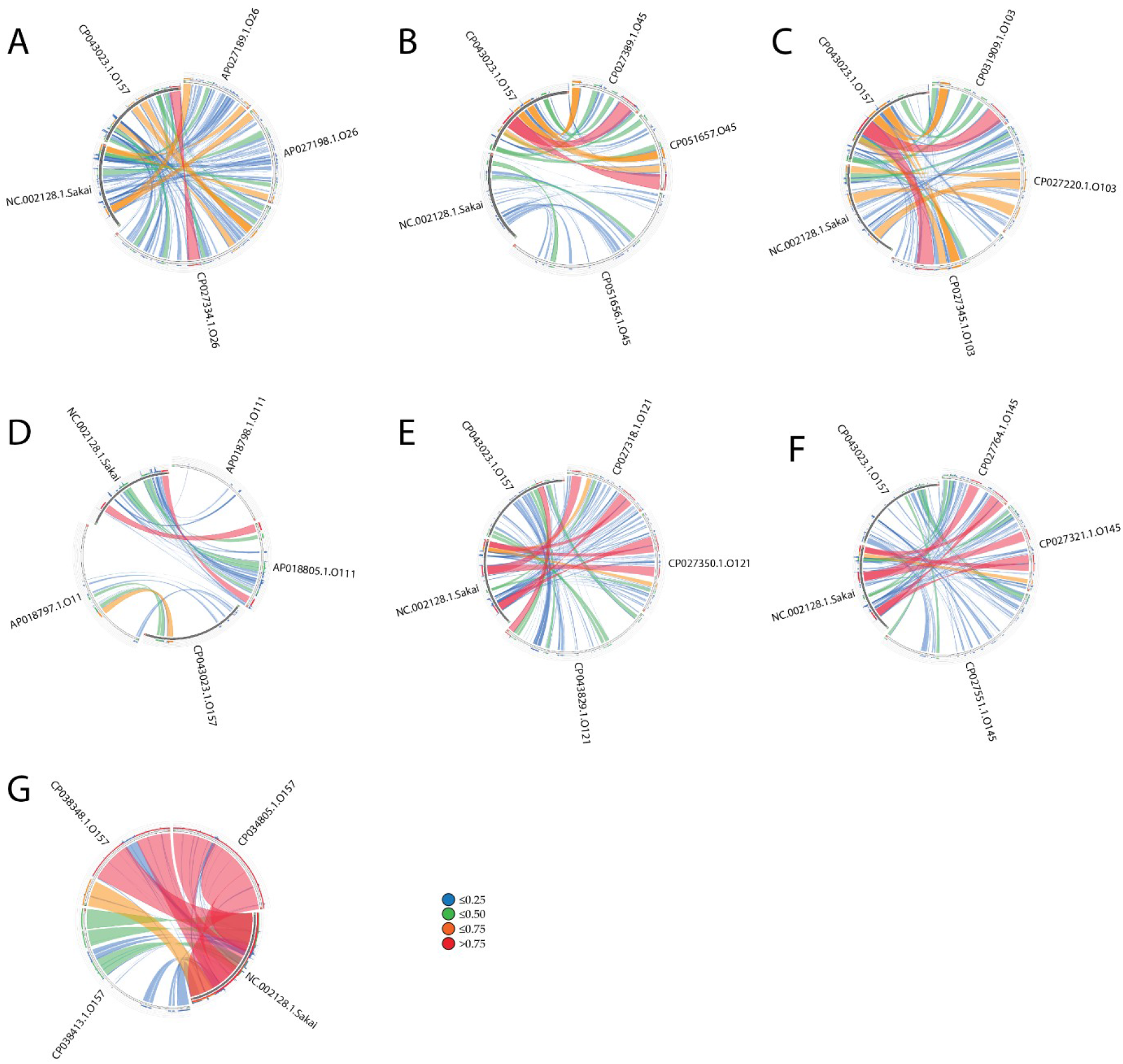
Circoletto-based pairwise comparison of STEC O157 plasmids with non-O157 serogroups. Panels (A–F) show O157 versus O26, O45, O103, O111, O121, and O145 plasmids, respectively. Panel (G) shows an internal comparison of the O157:H7 Sakai reference plasmid with O157 plasmids of different sizes. Ribbons represent shared genomic regions, with colors indicating sequence alignment scores: blue (≤0.25), green (>0.25 to 0.50), orange (>0.50 to 0.75), and red (>0.75).

### Enterohemolysin Operon across STEC Serogroups

Comparative alignment of the enterohemolysin (*ehx*) operon among *ehxA*-positive plasmids revealed a highly conserved gene organization across STEC serogroups (Fig. 4). Notably, among the 40 plasmids analyzed, only three showed structural differences compared to the others, highlighting the overall conservation of this locus (92.5%, 37/40). The operon, typically composed of *ehxC, ehxA, ehxB, and ehxD*, exhibited strong collinearity and high sequence similarity among plasmids belonging to O157, O26, O45, O103, O111, and O121 lineages. Although O26 and O103 strains are not closely related at the chromosomal level, the highest sequence conservation was observed among O26 and O103 plasmids, consistent with their close plasmid-level phylogenetic clustering (Fig. 5, Fig. 6). While the majority of plasmids harbor the canonical enterohemolysin operon, three plasmids displayed significant structural and sequence divergence. Phylogenetic reconstruction (Figs. S1–S4 in the Supplementary File) confirmed that these divergent operons belong to a distinct alpha-hemolysin clade, showing higher similarity to the *E. coli* 536 and pHly152 references than to the pO157-like group. Furthermore, we identified specific polymorphisms and pseudogenes, particularly in the *hlyB* and *hlyD* loci of certain O121 and O45 plasmids (e.g., CP043829.1 and CP051657.1; Figs. S1–S4 in the Supplementary File), suggesting a potential loss of hemolysin secretion capability in this specific plasmid backbone.

**Fig 4.**
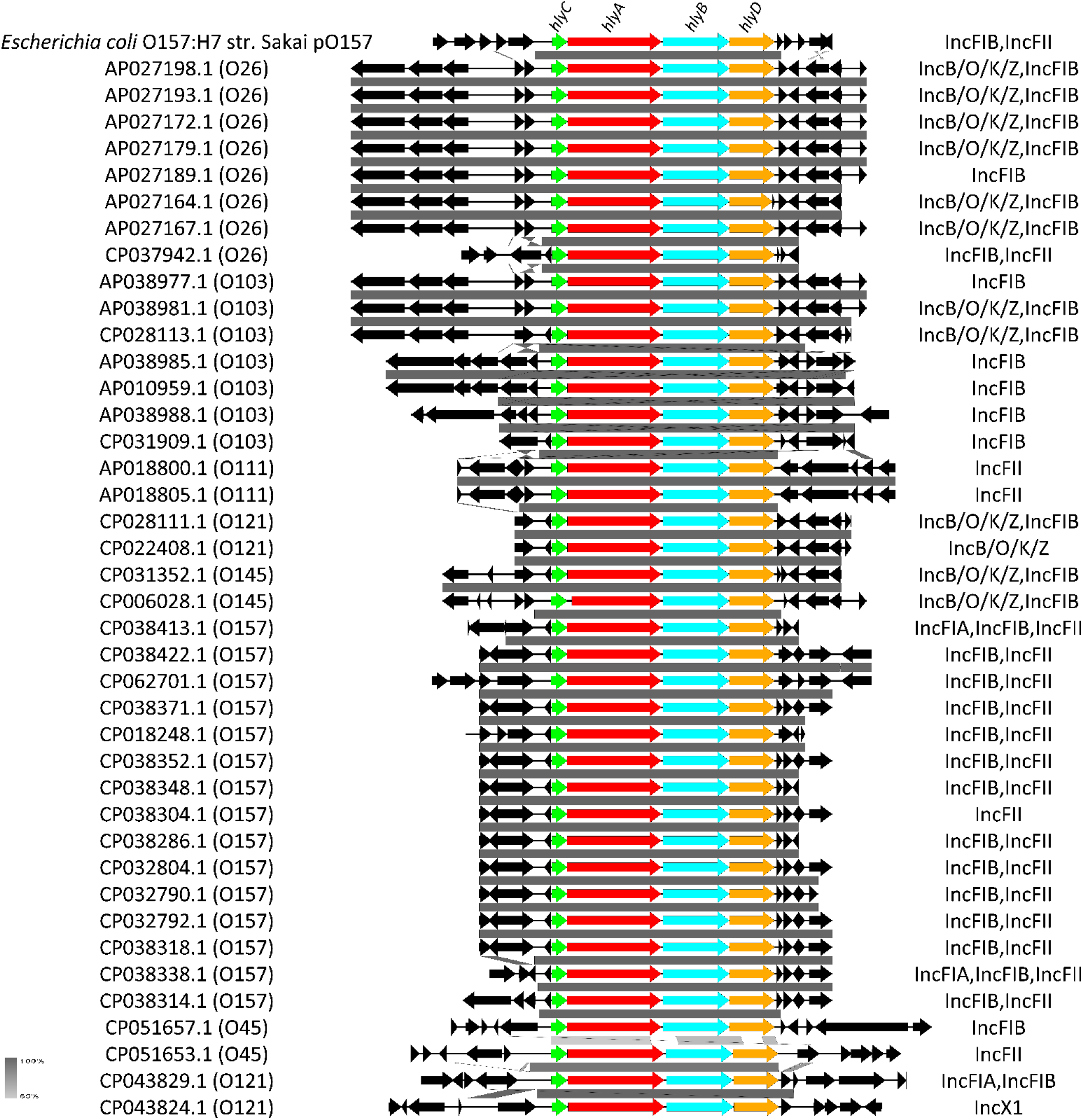
Comparative analysis of the genetic localization, organization, and protein homology of enterohemolysin operons among STEC plasmids, visualized using Easyfig.

**Fig 5.**
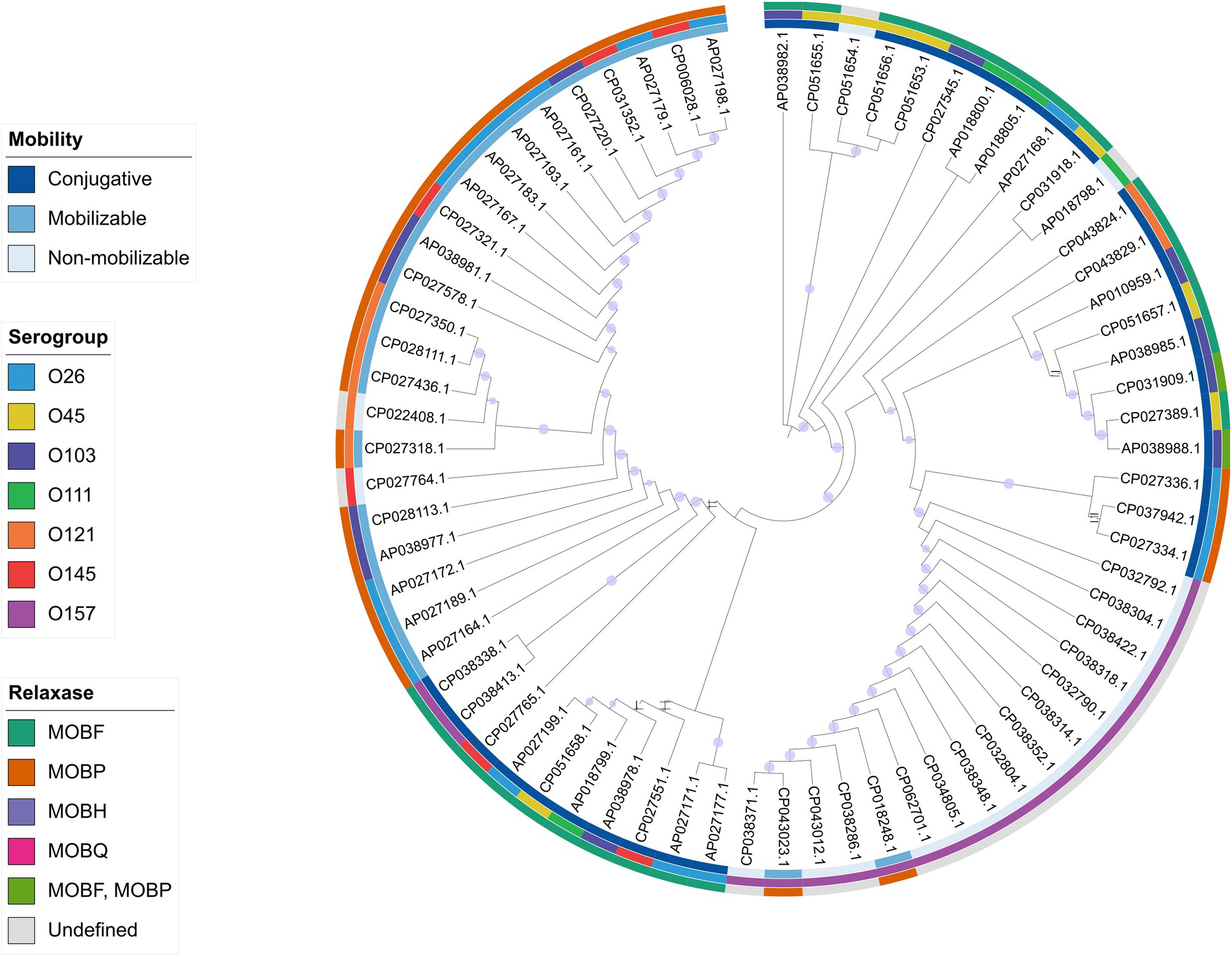
Maximum Likelihood phylogenetic tree of 71 STEC plasmids, inferred from concatenated *aidA*, *repB*, and *finO* sequences and displayed as a cladogram, showing clustering by O-group, relaxase type, and mobility, visualized in iTOL. Annotation rings are arranged from the innermost to the outermost layer as follows: serogroup, relaxase type, and mobility. Circles at internal nodes indicate adaptive bootstrap support ≥ 80%.

**Fig 6.**
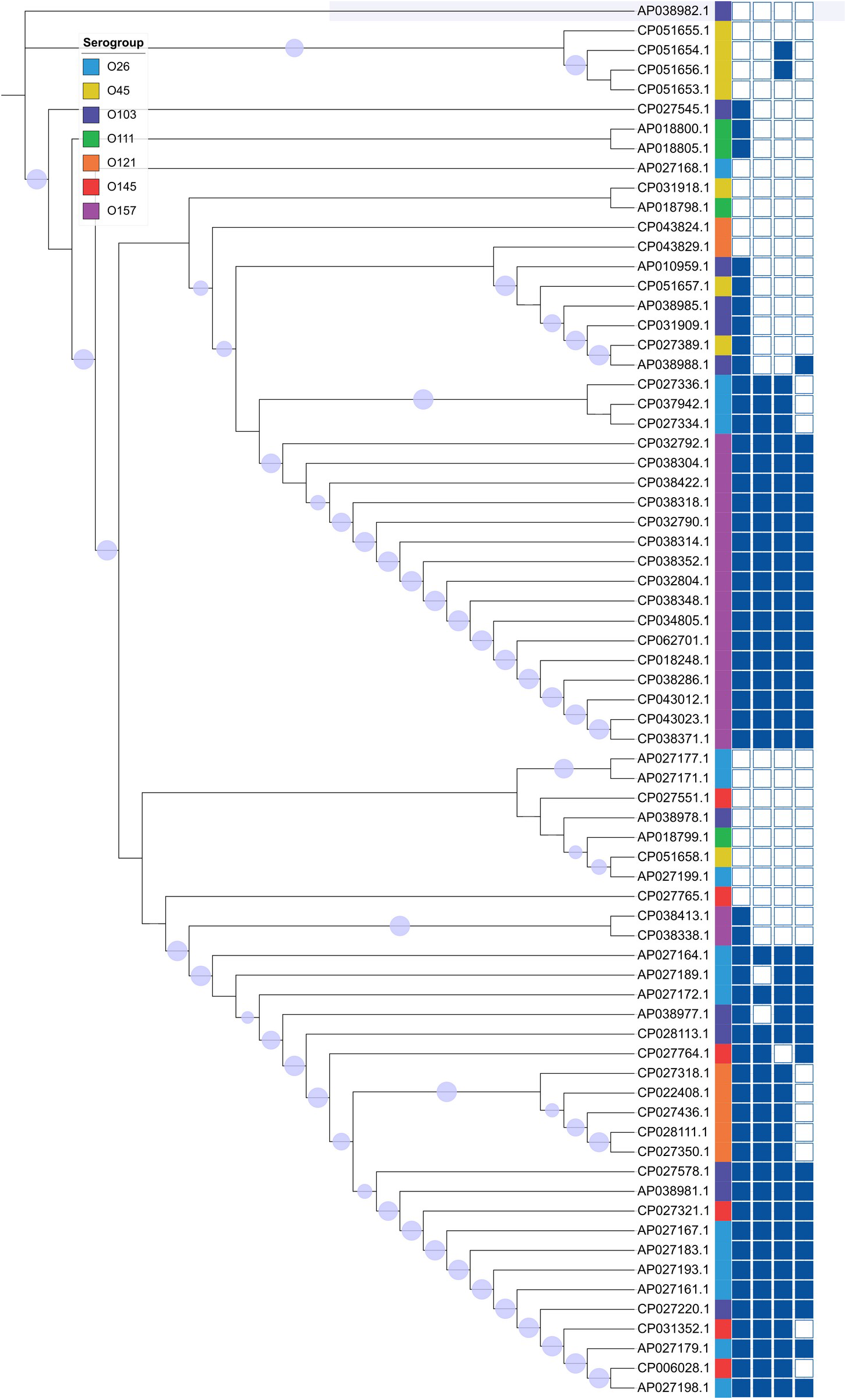
Maximum Likelihood phylogenetic tree of STEC plasmids constructed based on concatenated sequences of *aidA*, *repB*, and *finO* genes (same tree as Fig. 5), displayed as a cladogram and visualized using iTOL. Colored bars indicate plasmid serogroups, and the presence/absence of major virulence genes (*ehxA*, *toxB*, *espP*, and *katP*) is shown as a binary heatmap on the right. Circles at internal nodes indicate adaptive bootstrap support ≥ 80%.

### Phylogenetic Relationships and Clustering Patterns of Plasmids Based on Conserved Backbone Genes

The concatenated phylogenetic tree inferred from *aidA*, *repB*, and *finO* gene sequences revealed clear clustering patterns among the plasmids (Fig. 5). The central message of this analysis is that the plasmid backbone phylogeny recapitulates serogroup structure and co-varies with plasmid mobility: relaxase family, predicted mobility class and serogroup are not randomly distributed across the tree but are concordant along it, with O157 plasmids forming a single tightly clustered, predominantly non-mobilizable lineage and the non-O157 serogroups (notably O26/O103 and O45/O111) resolving as their own sub-clades. Plasmids belonging to the same serogroup, particularly O26, O121, and O157, tended to form several clusters throughout the tree, suggesting a possible evolutionary relationship and shared genetic background (Fig. 5). The topology of the tree was supported by high bootstrap values (median 99%; 45 of 68 internal nodes ≥ 90%), confirming the robustness of the inferred relationships. The relaxase typing data indicated that most plasmids carrying the MOBF family relaxase formed distinct clades. In contrast, MOBP and MOBF + MOBP plasmids appeared more dispersed across the tree, indicating horizontal transfer events (Fig. 5). Furthermore, plasmids with predicted conjugative potential were more broadly distributed across the phylogeny, reflecting their ability to transfer between diverse serogroups, whereas predicted mobilizable and predicted non-mobilizable plasmids were grouped into tighter clusters, suggesting a more limited host-range distribution across O-groups and reduced inter-lineage transfer (Fig. 5). Overall, the phylogenetic structure, supported by strong bootstrap values, suggests an association between plasmid backbone genes, mobility-related functions, and serogroup distribution.

Mapping the distribution of key virulence genes (*ehxA*, *toxB*, *espP*, and *katP*) onto the phylogenetic tree revealed distinct patterns of gene occurrence among plasmids from different O-groups (Fig. 6). These four genes were selected as they represent the most frequently detected plasmid-encoded virulence determinants in our dataset (42.9–68.8%) and are well-established contributors to STEC pathogenesis. The hemolysin gene *ehxA* was the most prevalent, widely distributed across nearly all major clades, particularly among plasmids associated with the O157 and O26 serogroups, indicating strong evolutionary conservation and an essential role in STEC virulence. In contrast, *toxB* was mainly restricted to O157 plasmids. The *espP* gene, encoding a serine protease autotransporter, showed a broader but uneven distribution, appearing in several O26 and O145 plasmids. Meanwhile, *katP*, responsible for catalase-peroxidase activity, was less frequent and primarily detected in O26, O103, and O157 plasmids (Fig. 6). Overall, this mapping underscores that the acquisition of virulence genes among STEC plasmids is both serogroup-dependent and lineage-specific, reinforcing the idea that plasmid-mediated virulence in STEC has evolved through a combination of vertical inheritance and horizontal gene transfer events.

### Chromosomal Phylogeny of STEC Strains Based on cgMLST

A total of 4,160 shared loci were identified, comprising 2,573 core and 1,587 accessory genes according to the inclusion and exclusion criteria of the SeqSphere+ Target Definer. While isolates belonging to the same serogroup frequently grouped together, several groups contained strains from different serogroups, indicating shared genomic backgrounds across distinct O-groups (Fig. 7). Notably, O157 strains, predominantly belonging to sequence type 11 (ST11), formed a relatively compact and well-defined lineage characterized by low allelic distances, reflecting strong genomic relatedness.

**Fig 7.**
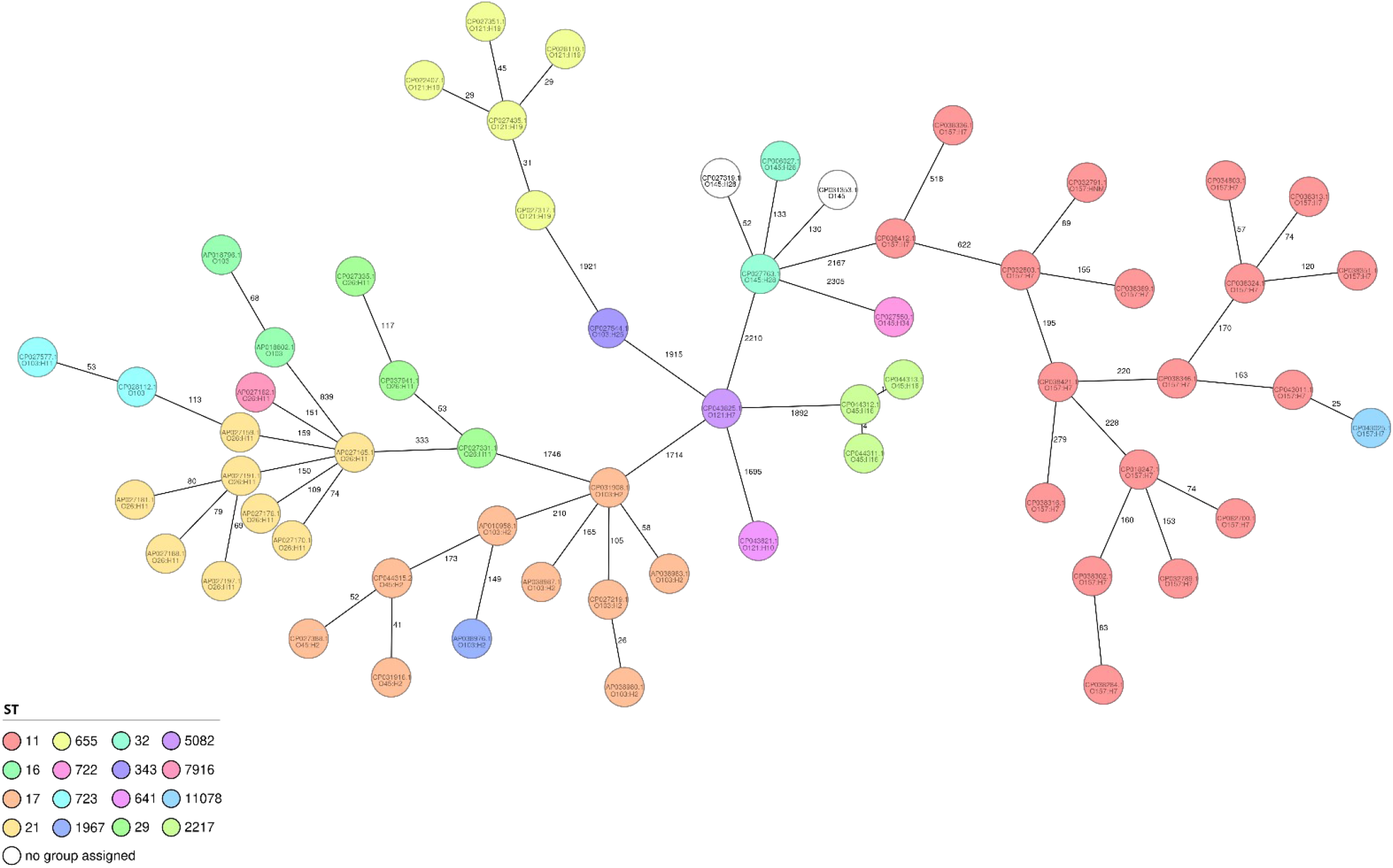
Chromosomal relatedness among the “Top Seven” STEC strains from which the analyzed plasmids were derived was determined using cgMLST in Ridom SeqSphere+ with the closed chromosome of *E. coli* K-12 substrain MG1655 as the seed. The strains’ Sequence Type (ST) was determined using the seven-gene *E. coli* MLST scheme available through EnteroBase. Numbers on connecting branches indicate the number of genome loci with differing allele status.

## Discussion

This study provides in-depth insight into the diversity, mobility, and virulence composition of STEC plasmids. By combining phylogenetic and comparative genomic approaches, we identified notable associations between plasmid backbone genes, replicon and relaxase types, and the distribution of key virulence factors. Importantly, isolates within each serogroup (O) represent diverse serotypes (O:H) and genetic backgrounds (see Supplementary File), so the observed associations between plasmid content and serogroups should be interpreted as trends rather than strict serogroup-specific features.

### Diversity and Distribution of Plasmid Replicon Types among “Top Seven” STEC Serogroups

Analysis of the plasmid replicons revealed distinct but diverse patterns among the “Top Seven” STEC serogroups. The IncFIB and IncFII replicons predominated, frequently co-localized on the same plasmids. F-type plasmids are well known for their stability and ability to exist in different *E. coli* pathotypes, which aids dissemination and persistence in bacterial populations (5, 52). The broad distribution of IncFIB across several serogroups also highlights its important role in carrying virulence genes. The frequent association of IncF with other replicons, such as IncB/O/K/Z, suggests that these plasmids have undergone recombination and adaptation over time (5, 53). Importantly, these hybrid IncB/O/K/Z+IncF-like plasmids are frequently found in non-O157 serogroups, introducing highly diverse genetic backbones. This variation shows that plasmid evolution in non-O157 strains is highly flexible and does not always follow the uniform, predictable patterns seen in O157 lineages. Overall, the results suggest that plasmids with stable backbones, such as those of the IncF family, are favored during STEC evolution because they support both survival and enable gene exchange between strains. The replicon architecture separates the serogroups in a way that parallels their chromosomal relatedness (Fig. 7). O26, O103, O121 and O145 plasmids frequently combined IncFIB with IncB/O/K/Z (8/18, 4/12, 5/7 and 4/8, respectively), whereas O45 plasmids lacked this combination entirely and carried IncFIB alone or with IncFII, O111 plasmids carried IncFII or IncB/O/K/Z+IncQ1, and O157 plasmids were restricted to IncF multireplicon backbones (IncFIB+IncFII or IncFIA+IncFIB+IncFII, 18/19) with no IncB/O/K/Z replicon. IncB/O/K/Z replicons occur across several Enterobacterales genera and are frequently associated with conjugative transfer and antimicrobial resistance (54, 55), so their presence in the non-O157 lineages may reflect access to a broader accessory-gene pool than that of the lineage-restricted IncF plasmids of O157. Consistent with this, the comparatively uniform IncF backbones of O157 coincided with limited predicted mobility (14/19 non-mobilizable) and with resistance genes in only 1 of 19 plasmids.

### Associations Between Replicon Types, Virulence Genes, and Plasmid Mobility

Clear associations between plasmid replicon types and major plasmid-associated traits such as virulence genes, relaxase families, and mobility have been observed (Fig. 1). Among them, F-type plasmids (IncFIB and IncFII) were most frequently associated with the key virulence genes *ehxA*, *toxB*, *espP*, and *katP*. F-type plasmids are therefore the main vehicles of these virulence genes in the collection; whether their replication stability and conjugative potential were selected together with pathogenicity cannot be inferred from sequence data alone (56). However, while this classic pattern is highly conserved in O157, the IncF replicons in non-O157 serogroups often co-occurred as hybrids with IncB/O/K/Z elements. These hybrid arrangements can alter standard virulence and mobility combinations. Overall, these patterns indicate a close relationship between plasmid replication, transfer ability, and virulence, facilitating the successful dissemination of F-type and hybrid plasmids among “Top Seven” STEC strains (57).

### Plasmid-Encoded Virulence Gene Profiles Reflect Lineage-Specific Adaptation in STEC

The distribution of plasmid-encoded virulence genes among the “Top Seven” STEC serogroups revealed clear lineage-associated patterns, suggesting possible adaptation to distinct host niches and infection strategies. Despite this diversity, *ehxA* emerged as the most consistently detected virulence determinant, supporting its central role in hemolytic activity and host cell damage across STEC lineages (58). This module was confined to the cargo-carrying plasmids of Table S2; the 32 plasmids without cargo carried at most colicin determinants, so the lineage-specific virulence signal described below resides in a defined subset of the STEC plasmidome.

In O26 and O103 plasmids, *ehxA*, *espP*, *katP*, and *toxB* frequently co-occur. Based on the individually characterized roles of these genes in hemolysis, proteolysis, and adherence (59), their combined presence may plausibly contribute to intestinal colonization and epithelial disruption (7), although the functional synergy among these gene products warrants further investigation. Supporting this, the PTU analysis classified these plasmids within closely related taxonomic units, indicating a stable evolutionary lineage dedicated to virulence maintenance. Furthermore, Circoletto-based alignments provided a sequence-level validation of this relationship, revealing extensive regions of high sequence similarity and conserved synteny between O157 and both O26 and O103 plasmids. In particular, the strong conservation of the core backbone observed among O157 plasmids (Fig. 3, G) reflects their shared PTU-E5 ancestry, a finding further corroborated by the whole-plasmid alignment comparison. The PTU-E5-associated O157 plasmids, along with the closely related O26 and O103 plasmids (both PTU-E69), carry the *ehxA*, *espP*, *katP*, and *toxB* module, representing a stably maintained, multifactorial virulence profile. This distinct combination of PTU and virulence gene content could be used for risk assessment, helping prioritize isolates during clinical or food-safety screening.

Plasmids from O45 and O111 strains were predominantly characterized by genes encoding the Type IV transfer apparatus, including the transfer-associated genes *traT* and *traJ*. These were often co-occurring with *anr* and *etpD*, a gene set associated with adhesion, secretion, and serum resistance (60, 61). This combination implies that these plasmids have diversified toward mechanisms that promote persistence in the host and protection against immune defenses, reflecting functional versatility rather than simple virulence attenuation. O45 plasmids were, in addition, the only ones to carry hlyA (3/8), cnf2, iha, and the afaA–D and cdt-IIIB determinants (1–2 plasmids each), an alternative, extraintestinal-type repertoire; *anr* and *etpD*, by contrast, were no more frequent in O45 than elsewhere. In O111, traT (5/5) rather than ehxA (2/5) was the most consistently detected determinant. Consistent with the presence of *traT* and *traJ* transfer genes, O45 and O111 serogroups also harbored the highest proportion of plasmids with predicted conjugative potential, suggesting a strong potential for horizontal gene transfer.

In O121 and O145 serogroups, the predominance of *ehxA*, *espP*, and *toxB* indicates conservation of a virulence module similar to that of O26, suggesting possible convergent evolution among non-O157 STEC toward shared colonization and toxin-associated pathways.

Conversely, O157 plasmids exhibited the most complex virulence architecture, typically integrating *anr*, *ehxA*, *espP*, *etpD*, *katP*, and *toxB*. This combination represents an expanded virulence network capable of mediating multiple infection stages, from adhesion and toxin delivery to immune evasion. Such a configuration is consistent with the classical pO157 reported in highly virulent O157:H7 strains, suggesting a recurrent co-occurrence of plasmid-encoded virulence functions within this lineage (62). Among all serogroups, *etpD* was strongly associated with O157, detected in 94.7% (18/19) of O157 plasmids versus 8 of 90 non-O157 plasmids (Fisher’s exact test, P = 1.2 × 10⁻⁷).

### Limited occurrence and distribution of antimicrobial resistance genes among STEC plasmids

The investigation revealed a relatively low prevalence of antimicrobial resistance (AMR) genes among the analyzed STEC plasmids, with approximately 11% (12/109) encoding resistance determinants, consistent with findings from several large surveillance studies (53, 63). This observation suggests that virulence, rather than antimicrobial resistance, remains the predominant adaptive strategy within the “Top Seven” STEC plasmidome. AMR-positive plasmids were mainly identified in the O26 and O111 serogroups; 10 of the 12 were classified as having predicted conjugative potential and 9 encoded the virulence gene *traT*. The *traT* gene is typically located within the transfer (*tra*) region of plasmids with predicted conjugative potential and is associated with the transfer machinery, while also contributing to serum resistance by expressing an outer membrane protein (64). Its prevalence in MDR *E. coli* strains has also been reported in previous studies (65).

Although only a small proportion of STEC plasmids (∼11%) carried antimicrobial resistance genes, most of these AMR-positive plasmids (92%, 11/12) encoded resistance to multiple antibiotic classes, indicating a relatively high level of MDR among the resistant subset. The diversity of detected AMR genes, including *aph(3’)-Ia*, *sul1*, *tet(A)*, and *bla*TEM-1B, reflects resistance mainly to aminoglycosides, sulfonamides, tetracyclines, and beta-lactams, respectively. This suggests that, while AMR is uncommon in STEC plasmids overall, when resistance is present it is usually found in combination with other resistance genes rather than as a single trait.

Despite this, the overall low frequency of AMR-positive plasmids clearly distinguishes STEC from extraintestinal pathogenic *E. coli* (ExPEC), which commonly harbor highly resistant plasmids (66). The occasional detection of clinically important resistance genes such as *mcr-5.1* and *mph(A)*, particularly in O111 plasmids, is concerning, as these genes confer resistance to last-resort antibiotics and indicate sporadic horizontal gene transfer from other Enterobacterales.

Although O157 plasmids encoded the highest number of virulence genes, they harbored few AMR determinants. This pattern may indicate retention of virulence- and fitness-associated functions rather than acquiring AMR determinants (67). This pattern aligns with previous observations that highly virulent STEC serotypes, particularly O157:H7, rarely harbor resistance genes compared to commensal or ExPEC strains (53, 63). Altogether, these findings indicate that while AMR remains limited in STEC plasmids, its occurrence is frequently linked to multidrug resistance, underscoring the importance of continued genomic surveillance to monitor the emergence and spread of MDR plasmids within STEC populations.

Interestingly, *lpxM* and *eptC* were identified in 48.6% (53/109) and 40.4% (44/109) of the 109 plasmids, respectively (Table S4). While these genes were also present on the chromosomes of some isolates, the plasmid-borne variants exhibited distinct nucleotide and protein sequence differences. We hypothesize that these plasmid-derived homologs may contribute to reduced susceptibility against membrane-disrupting antibiotics, such as colistin and polymyxin B, by modifying the lipid A moiety of the lipopolysaccharide (LPS). The observed sequence variations from chromosomal counterparts might affect their identification by standard genomic databases, potentially leading to an underestimation of the plasmid-mediated resistome in STEC. Further functional studies are required to determine whether these sequence differences impact the efficacy of membrane-targeting antimicrobial agents.

### Relaxase Types and the Evolutionary Implications of Plasmid Mobility

The relaxase typing results revealed considerable diversity among the analyzed STEC plasmids, illustrating how different plasmid families contribute to horizontal gene transfer and bacterial adaptation. The predominance of MOBP and MOBF relaxases highlights two major mobility systems that shape plasmid evolution within STEC populations. MOBF-type plasmids were invariably linked with predicted conjugative elements capable of autonomous transfer between strains. In contrast, MOBP-type plasmids were commonly associated with predicted mobilizable plasmids that rely on conjugative partners for transfer (34). This pattern suggests that MOBF plasmids may play a central role in disseminating mobile genetic elements both within and across STEC lineages.

Interestingly, the highest proportion of plasmids with predicted conjugative potential was detected in the O45 and O111 serogroups, while O157 plasmids were predominantly predicted non-mobilizable. This pattern may be consistent with evolutionary trade-offs, in which O157 plasmids have accumulated virulence and maintenance genes at the expense of mobility (68). However, experimental validation would be required to confirm this hypothesis, as plasmid mobility was assessed *in silico*.

A notable proportion of the cargo-carrying plasmids in this study (26.0%, 20/77; 14 of them from O157) lacked a defined relaxase type, possibly due to novel or highly divergent mobility systems that remain uncharacterized in current databases, or to deletions and rearrangements within relaxase-coding regions (69). Alternatively, a subset of these plasmids may rely on helper plasmids for mobilization, suggesting a coexistence strategy based on inter-plasmid dependency (70). This observation underscores the extensive genetic diversity and evolutionary flexibility of STEC plasmids beyond the conventional MOB family classification. Overall, these findings indicate that plasmid mobility in STEC is not random but structured, shaped by evolutionary pressures that balance stability, transfer potential, and the maintenance of virulence-associated genes. The observed relaxase diversity likely represents ongoing evolutionary adaptation to both host and environmental contexts.

The 32 plasmids without virulence or resistance cargo complement this picture. Their gene content was restricted at most to colicin determinants, and the pO157-type virulence module was absent, whereas every one of them received a predicted mobility class, with conjugative machinery confined to plasmids above 37 kb and relaxase-only (mobilizable) plasmids dominating below 11 kb; across the whole collection no conjugative plasmid was smaller than 20.8 kb. This size–mobility partition is consistent with the general relationship between plasmid size and self-transmissibility described across bacterial plasmids (68), and indicates that the smallest STEC plasmids behave as hitchhikers that depend on co-resident conjugative plasmids, including the large IncF virulence plasmids described above, for their dissemination. Because all 109 plasmids were processed through one pipeline at one documented tool version, the two cargo classes can be compared directly. We note that nearest-neighbor distances against the MOB-suite reference database cannot serve as independent corroboration for these plasmids, as they are already contained in that database.

### Plasmid Sequence Typing and Evolutionary Lineages in STEC

The IncF subclassification results further clarified the evolutionary structuring of STEC plasmids. The F-:A-:B11 sequence type was the most prevalent and widely distributed across multiple serogroups, including O26, O103, O121, and O145, suggesting the presence of a conserved plasmid lineage circulating among diverse STEC populations (26). Its presence across unrelated lineages indicates a successful plasmid backbone that may facilitate the stable maintenance of virulence determinants such as *ehxA*, *toxB*, and *espP*. Notably, the majority of plasmids assigned to this sequence type (95.5%, 21/22) were classified as predicted mobilizable and may disseminate via helper plasmids. In contrast, sequence types F13:A-:B-, F5:A-:B-, and F23:A-:B3 were restricted to O45, O111, and O157 plasmids, respectively, pointing to lineage-specific plasmid evolution and potential adaptation to host or ecological niches. Consistent with PTU assignments, most O157 plasmids clustered within a single dominant PTU (PTU-E5), with only one plasmid assigned to PTU-C. In contrast, plasmids from O26, O121, O145, and a subset of O103 isolates were predominantly associated with PTU-E69, reflecting shared plasmid backbones across these serogroups. Only minor discrepancies were observed, mainly involving a few singleton PTU-? assignments, further supporting the overall concordance between PTU classification, pMLST typing, and plasmid phylogenetic relationships shown in Fig. 6. PTU-E69 was almost exclusively an IncFIB+IncB/O/K/Z lineage: 20 of its 23 members carried this replicon combination, distributed across O26, O103, O121 and O145, whereas O45 plasmids lacked the combination entirely and fell into PTU-E5, PTU-FE and PTU-E77, and O111 plasmids into PTU-E41 and PTU-B/O/K/Z. The concordance between PTU, replicon architecture and serogroup therefore extends the chromosomal lineage structure (Fig. 7) to the plasmid level.

### Conservation of the Enterohemolysin Operon among STEC Plasmids

EHEC-associated enterohemolysin (*ehx*) was acquired through independent acquisition of virulence plasmids by distinct O157 and non-O157 lineages (8). The comparative genomic assessment of *ehxA*-positive STEC plasmids highlighted a remarkably conserved locus across the major “Top Seven” serogroups. Rather than showing extensive structural divergence, the operon preserved a stable genetic framework consisting of *ehxC, ehxA, ehxB, and ehxD*, indicating that this virulence module has been maintained with little genomic rearrangement over time (8).

Slight sequence variations were observed only in short intergenic or adjacent hypothetical regions, showing minor differences between lineages while the operon’s main function remained conserved (58, 71). Notably, plasmids belonging to the O26 and O103 serogroups showed the highest degree of sequence similarity, reflecting close evolutionary relatedness at the plasmid level and potentially similar mechanisms of virulence expression (72). These findings are consistent with the plasmid-encoded virulence gene profiles and plasmid-level phylogenetic clustering observed for O26 and O103, further supporting their close relationship at the plasmid level.

Furthermore, we identified specific polymorphisms and pseudogenes, particularly in the *hlyB* and *hlyD* loci of certain O121 and O45 plasmids. Consistent with previous reports (73), these alterations, such as the truncated *hlyB* found in some lineages, may indicate reduced or absent hemolytic activity. This suggests that the sporadic occurrence of alpha-hemolysin variants and localized genetic degradation (pseudogenization) contribute to the functional and evolutionary diversity of hemolysin-positive STEC plasmids.

The strong conservation of the *ehx* operon across genetically distinct plasmids suggests that selective forces favor its retention, likely due to its roles in hemolysis and host interaction. Collectively, these findings indicate that the *ehx* operon serves as an evolutionarily stable and functionally significant virulence determinant in STEC plasmids, representing a core element of their pathogenic identity.

### Correlation Between cgMLST Clades and Plasmid Diversity

Although strains belonging to the same serogroup often grouped together, several clades included isolates from different O-groups, indicating shared genomic backgrounds beyond serogroup acquisition. Notably, the consistent similarity in the genetic framework of O157 aligns with a relatively conserved and virulence-associated plasmid profile, suggesting a tight co-evolution between chromosomal and plasmid backgrounds in O157 lineages. In contrast, the broader diversity of the chromosomal backbone observed among non-O157 strains parallels the higher variability in their plasmid replicon types, gene content, mobility, and virulence repertoires. This pattern supports the idea that STEC plasmids evolve through a combination of vertical inheritance within lineages and horizontal transfer across strains, allowing key virulence modules to spread independently of the genetic makeup (74).

Although the STEC strains in this study were collected from diverse sources, countries, and over a broad time span (2016–2025), no relationship was observed between STEC plasmid profiles and the source, year, or location of isolation. This suggests that these STEC-associated plasmids are relatively stable over time and widely distributed across different hosts and regions. It indicates that they can be transferred between strains from different origins, contributing to the dissemination of virulence and resistance genes.

This analysis is based on a snapshot of the publicly available plasmid inventory of the seven major STEC serogroups in NCBI at the time of retrieval (July 2025). The functional consequences of the identified gene combinations and mobility predictions were inferred from sequence data alone and remain to be validated experimentally.

## Conclusions

This study provides a comprehensive overview of the diversity and evolutionary patterns of plasmids in the “Top Seven” STEC serogroups. Our findings indicate that these plasmids exhibit structured genetic architectures combining virulence-associated genes with stability and mobility elements. The co-occurrence of conserved backbones and lineage-specific features points to ongoing plasmid-mediated adaptation and horizontal exchange across STEC lineages, alongside the influence of stochastic processes and genetic drift.

Overall, our data support the view that plasmids contribute to the maintenance and dissemination of virulence traits across different STEC lineages and ecological contexts. Notably, plasmids from the O26 and O103 serogroups exhibited a close relationship, consistent with their shared plasmid-level phylogenetic grouping and relatively high-risk virulence gene profiles, although clinical disease outcomes are ultimately determined by a complex interplay of factors, including host age, immune status, and infecting dose. In contrast, the O121/O145 group was distinguished by a moderate virulence gene repertoire, whereas plasmids from O45 and O111 appeared distinct, particularly in their transfer-associated gene content. Finally, O157 plasmids harbored the highest number of virulence-associated genes and formed a distinct clade, consistent with the virulence plasmid repertoire described for this lineage.

## List of abbreviations

AMR: Antimicrobial Resistance
STEC: Shiga toxin-producing *Escherichia coli*
NCBI: National Center for Biotechnology Information
HUS: Hemolytic Uremic Syndrome
PTU: Plasmid Taxonomic Unit
CARD: Comprehensive Antibiotic Resistance Database
pMLST: Plasmid Multilocus Sequence Typing
BLAST: Basic Local Alignment Search Tool
MAFFT: Multiple Alignment using Fast Fourier Transform
MEGA: Molecular Evolutionary Genetics Analysis
iTOL: Interactive Tree of Life
cgMLST: Core Genome Multilocus Sequence Typing
MDR: Multidrug-Resistant

## Declarations

### Ethics approval and consent to participate

This study did not involve any experiments on humans or animals. All genomic data analyzed were obtained from NCBI. Therefore, ethical approval and informed consent were not required.

### Consent for publication

All authors have read and approved the final version of the manuscript and consent to its publication.

### Availability of data and materials

All data supporting the findings of this study are included in the manuscript and its supplementary materials. The accession numbers for all whole-genome sequences analyzed in this study are provided in the article and the supplementary files. Additional information, including FASTA sequences of the analyzed plasmids and chromosomes as well as raw output files generated from the genomic analyses, can be obtained from the corresponding author upon reasonable request. The PlasmidTyper suite used to submit, retrieve and collate the plasmid typing results, together with the accession list and compiled profiles of the 109-plasmid panel, is available at https://github.com/kramppe/PlasmidTyper (v1.0.1).

### Authors’ contributions

A.N. and M.E. designed the study, supervised the project, reviewed the data analyses, and drafted the manuscript. M.E., I.R., S.M., A.D., N.D., S.S.K.K., A.A., A.K., A.S.M., and A.T.S. conducted the genomics analyses. M.E. and S.S.K.K. developed the PlasmidTyper software used for plasmid taxonomy, mobility, and gene-inventory typing. U.R. and F.G. reviewed the data analyses and edited the manuscript. All authors have read and agreed to the published version of the manuscript.

## Acknowledgment

The authors acknowledge the Research Computing Support Group (RCSG) at UT San Antonio for providing computational and high-performance computing resources on the Advanced Research Computing (ARC) cluster that contributed to the reported research. The authors would like to thank Mr. Hamed Nemati for his valuable assistance in figure design. We also sincerely appreciate Ms. Nazanin Shadanpour from the Research Institute of Biotechnology, Ferdowsi University of Mashhad, for her insightful feedback and constructive comments during this study. We thank Jacob Alford for assistance with plasmid profiling.

